# Inhibition of the Adenosine pathway activates the immune response against Mesothelioma

**DOI:** 10.64898/2026.05.08.722957

**Authors:** Caterina Costa, Steven G. Gray, Giulia Pinton, Laura Moro, Erika Del Grosso, Cristiana Bellan, Laura Addi, Rita Lombardi, Francesca Bruzzese, Davide De Biase, Biagio Pucci, Elena Di Gennaro, Paolo Antonio Ascierto, Giovanni Luca Gravina, Luciano Mutti

## Abstract

**Background:** Mesothelioma (Me) is an aggressive cancer with limited response to conventional therapies. The tumor’s harsh microenvironment contributes to immune escape and therapy resistance and the effects of ICIs on Me are still unclear. Adenosine, an immunosuppressive molecule produced from AMP by the enzyme CD73, accumulates in hypoxic tumor areas. Elevated CD73 and adenosine receptor A2B (A2Br) levels on Me cells are linked to worse patient outcomes, indicating their important role in disease progression and potential as targets for treatment.

**Aim:** This study characterizes the Me-ME (micro environment) and evaluates the efficacy of TT-4 (A2B inibitor) and AB680 (CD73 inibitor), alone or with aPD-1, using 3D models *in vitro* and *in vivo*.

**Methods:** CD73 and A2B receptor levels were quantified in tumor and normal samples using qRT-PCR and IHC. Cells lines were treated with CoCl_2_ to mimic hypoxia, then CD73, A2Br and related markers were analyzed. MSTO-211H and REN cells were silenced for CD73, grown as spheroids and adenosine release was measured. Co-culture spheroids of MSTO-211H and Jurkat cells were treated with AMP and CD73 inhibitor, then analyzed for viability and immune markers. An orthotopic Me model was established by injecting AB1-B/c-LUC cells and monitored by *in vivo* imaging. Proteomic analysis of spheroids was conducted to identify proteins and pathways involved.

**Results:** Hypoxia boosts CD73 and A2Br expression in Me cells, leading to adenosine production via CD73. In 3D co-cultures, AB680 lowered Me cell viability and enhanced activation of Jurkat T cells. In mice, combining aPD-1 therapy with A2Br or CD73 inhibitors strongly reduced tumor growth. Proteomics identified 93 proteins influenced by adenosine signaling through A2B.

**Conclusion:** Targeting the adenosine pathway alongside PD-1 blockade offers a promising new immunotherapy strategy for Me.

## Introduction

Mesothelioma (Me) is a highly aggressive malignancy that arises from mesothelial-lined surfaces, such as the pleura or the peritoneum, and is related to the environmental or working exposure to asbestos ^1, 2^ Although the industrial use of asbestos has been banned in Western industrialized the incidence of Me is expected to continue to increase for the next few decades due to the past widespread usage of these fibers ^3, 4^. Treatment of Me is challenging for different reasons, including the fact that most of the cases are diagnosed at an advanced stage and treated with palliative systemic chemotherapy ^5^. Although Immune-checkpoint inhibitors (ICIs) for human cancer have positively impacted the treatment of some tumors, the evidence suggesting their actual clinical relevance in Me has been queried by several groups ^6-8^. In addition, patient stratification based on a deeper insight of the immune response to cancer is urgently needed ^9^. Despite initial observations suggesting the use of ICIs for ME ^10^ ME has shown poor response to the most updated clinical trials including immunotherapy ^11 12 13^ and an analysis of the recently approved combo Ipilimumab/Nivolumab ^14^; ^15^ shows many pivotal critical points cannot be ignored ^8, 16^. Moreover, recent RWE studies ^17-19^ along with a series of negative trials ^13, 20^ provided additional evidence of the need for new approaches for immunotherapy for this stubborn neoplasm. It seems clear that the poor performance of these treatments is largely due to the need of a deeper specific understanding of the complex ME immune-biology ^21^.

Many other biological factors may play a role in the failure of ICIs in human tumors: known issues include acidic and hypoxic tumor microenvironments, lymphocytes exclusion and Cancer Associated Fibroblasts (CAFs) ^12, 22^. Hypoxia has been shown to promote the activation of the adenosine pathway including the key ectoenzymes CD39 and CD73 (also known as nt5E) ^23-26^ that degrade ATP to AMP and AMP to Adenosine respectively. whose signaling, in turn, is mediated by ADORA2A and -B receptors (A2Ar and A2Br) ^27^.

Adenosine binding with these receptors on tumor infiltrating inflammatory cells leads to the iMediate inactivation of tumor specific CTLs and other well recognized immune effectors (*Ref*) and is able to induce recruitment and changes of the genomic profile of immune-suppressive regulatory T cell (Tregs) subpopulations (Oliveto et al., manuscript in preparation). Thus inhibitors targeting this pathway (are progressively attracting the attention as actionable targets for immunotherapy in many solid tumors alone or in combination with ICI ^28-34^. In particular hypoxic tumors like Me ^35^ could activate the adenosine receptors-related pathway, providing a major immunosuppressive mechanism ^36^, CD73/adenosine receptors expression and adenosine-related gene-expression signature have been evoked to select patients for immunotherapy with adenosine inhibitors ^30, 37^. Our results complement in silico analyses of existing datasets and reveal an inverse correlation between A2Br and CD73 expression in patients with Me. Because both CD73 and A2B receptor are actionable targets with compounds already available in clinical settings, the purinergic pathway offers the rationale for being tested as a novel specific immune therapy for Me.

## Materials and methods

### Reagents and Antibodies

59-(N-Ethylcarboxamido) adenosine (NECA, ab120440), a general adenosine receptor agonist that targets all subtypes (ADORA1, 2A, 2B and 3) and MRS 1754 [N-(4-cyanophenyl)-2-[4-(2,3,6,7-tetrahydro-2,6-dioxo-1,3-dipropyl-1H-purin-8-yl)-phenoxy]acetamide] (M6316), a specific A2B receptors antagonist (A2Br) were purchased from Abcam and SigmaAldrich, respectively. Cobalt dichloride (CoCL2, 232696; SigmaAldrich), a chemical inducer of Hypoxia-Inducible Factor-1 alpha (HIF-1α and hypoxia). Adenosine was sourced from Sigma-Aldrich (Cat. No. A9251). LC-MS grade formic acid (FA), methanol (MeOH) and water were purchased from Carlo Erba (Milan, Italy, Cat. 414831 and 412111). Specific antibodies used for Western blot and IHC analysis were as follows: rabbit polyclonal anti-HIF-1α (3716S, Cell Signaling), anti-CD73 (ab133582, Abcam), anti-A2Br (ab229671, Abcam) and anti-GAPDH (14C10-2118L; Cell Signaling Technologies).

### Case selection

Formalin-fixed and paraffin-embedded (FFPE) samples were obtained at the archives of the Siena University Hospital (AOUS) and collected to evaluate CD73 and A2Br expression by real-time qRT-PCR and immunohistochemistry (IHC). This is a retrospective mono-institutional analysis performed on 64 advanced ME patients’ diagnosis (of which 38 were epithelioid and 26 non-epithelioid) who received surgery and/or chemotherapy and/or radiotherapy. Mediastinal pleura removed during coronary artery bypass surgery was used as normal controls (7 control samples). Clinical records were reviewed to document sex, age of patient and histology of Me. For the immunohistochemistry (IHC) of human Me tissue, the same antibodies used for the Western blot analysis were employed, according to the manufacturer’s instructions.

### RNA Isolation and Real-time quantitative reverse transcription (qRT)-PCR

Total RNA was extracted from FFPE sections of tumor tissue and normal pleura samples using Recover All Total Nucleic Acid Isolation Kit for FFPE tissues (Ambion) following the manufacturer’s instructions. Starting from 500 ng of RNA, cDNA was synthesized using the iScript cDNA Synthesis kit (Bio-Rad). For *ADORA2B* and *nt5E* mRNA (encoding CD73 protein) expression analysis, cDNA was amplified in the 7900HT Fast Real-Time PCR using the Power Sybr Green Mix (Applied Biosystems). Primer sequences are listed in **Supplementary Table S1**. Relative gene expression was calculated using the 2^−ΔΔct^ method relatively to controls (β-actin was used as housekeeping gene). Total RNA of multicellular spheroids was extracted using the guanidinium thiocyanate method. Starting from equal amounts of RNA, cDNA used as template for amplification was synthesized by the reverse transcription reaction using RevertAid Minus First Strand cDNA Synthesis Kit from Bio-Rad Laboratories S.r.l. (Segrate (MI), Italy), according to the manufacturer’s instructions. 50 ng of cDNA were used to perform RT-PCR amplification of mRNA.

### Cell Lines

Human mesothelioma cells were purchased from the American Type Culture Collection (ATCC), while murine cells were provided by Dr. Crippa Massimo ^41^, and were cultured under standard conditions in RPMI-1640 medium supplemented with 10% FBS, 1% L-glutamine and 1% penicillin-streptomycin at 37 ⍰C in humidified incubator with 5% CO2. The normal mesothelial NP2 cells developed at the University of Vienna ^38^ were provided by Dr. Steven Gray. All lines were then routinely tested for the presence of mycoplasma with the PlasmoTest ™ — Mycoplasma Detection Kit (Invivogen, Cat # rep-pt1).

### Cobalt chloride treatment

MSTO-211H and NCI-H28 cells were first cultured in complete growth medium in standard CO_2_ incubator. After 24 h, a 770 mM stock solution of cobalt chloride (CoCl_2_) was prepared in sterile distilled water and further diluted in RPMI to obtain the final desired concentrations (250 nM). Then, cell lines were incubated for 6 h with CoCl_2_ to mimic hypoxic conditions and determine hypoxia-associated genes by Western blotting (HIF-1α, CD73 and A2Br) and qRT-PCR (*ADORA2B* and *nt5E*) compared to untreated cells.

### Protein extraction, Western blotting

Analyses were carried out according to standard procedures. 50 μg of the total proteins were subjected to SDS-PAGE and then transferred to nitrocellulose membranes to evaluate HIF-1 α (as a control of hypoxia induction) CD73 and A2Br. After blocking (5% BSA for 30 min at room temperature), the membrane was incubated with primary antibodies in 5% BSA/PBS/Tween20 overnight at 4°C. Following 1 h at room temperature with the secondary antibodies, the immunodetection was performed using the Immobilon Forte Western HRP Substrate (Millipore).

### Multicellular Spheroids (MCSs)

The MSTO-211H and the REN cell line were grown to 80% confluence in tissue culture dishes and transiently transfected with negative control or specific *nt5E* siRNAs from Qiagen (Catalog No. 1027417 and 1027416 respectively Hilden, Germany) using LipofectAMINE transfection reagent (Life Technologies, Italy) as described by the manufacturer. After 24 hours cells were trypsinized, stained with Trypan blue and the number of cells considered viable was counted in a Burker chamber. Multicellular spheroids (MCSs) were generated in non-adsorbent round-bottomed 96-well plates. The 96-well plates were coated with a 1:24 dilution of polyHEMA (120 mg/ml) in 95% ethanol and dried at 37°C for 24 hours. Before use, the plates were sterilized by UV light for 30 minutes. For the generation of MCSs, 1×10^4^ cells/100 µl of RMPI 1640 medium were added to each well and culture for additional 48 hours. Spheroids were then incubated 1 hour with 50μM AMP and media were collected for semi-quantification analysis of adenosine.

### Adenosine semi-quantification via LC-MS/MS

The preparation of the adenosine stock solution (SS) involved two steps. First, a 5 mM water solution was prepared; then, a serial dilution with RPMI-1640 was carried out to obtain a set of 6 calibrators (0.1, 1.0, 2.5, 5.0, 7.5, 10.0 µM) for plotting a matrix-matched curve. This range covered the expected concentrations of the study samples. The samples were diluted 1:3 with MeOH (100 µL + 300 µL) to precipitate proteins, followed by a 1-minute vortexing and centrifugation at 13,000 rpm for 5 minutes.

Supernatant aliquots of 300 µl were transferred into 1.5 ml centrifuge micro-tubes and fully dried under vacuum at 30 °C for 4 hours. Subsequently, they were reconstituted with 80 µL of H2O 0.2% FA v/v before being analyzed using LC-MS/MS. Both the study samples and calibrators were prepared in a similar manner. The analyses were performed using the Q-Exactive Plus UHMR Hybrid Quadrupole Orbitrap™ Mass Spectrometer equipped with a Vanquish™ Duo UHPLC system (Waltham, MA, USA). The LC conditions included a Synergi 4 µm Polar-RP 80 Å, LC 150 × 2 mm column with a temperature of 45 °C.

Mobile phases consisted of A – H2O 0.1% FA and B – MeOH 0.1% FA. A flow rate of 0.300 mL/min was maintained, and gradient elution was achieved by changing the A-B ratio from 95:5, v/v to 5:95, v/v over 9.0 minutes, followed by A-B (5:95, v/v) until 11.0 minutes, and finally reaching A-B (95:5, v/v) at 11.5 minutes. The column was reconditioned with A-B (95:5, v/v) from 11.5 to 15.5 minutes, resulting in a total run time of 15.5 minutes. Each sample was injected with a volume of 5 µl. Regarding MS tuning, the ESI source was used with ESI+ ionization, setting the spray voltage at 3.50 kV and maintaining the capillary temperature at 300 °C. The sheath gas flow (N2) was set to 45.00 L/min, while the sweep gas flow (N2) and auxiliary gas flow (N2) were adjusted to 1.00 L/min and 10.00 L/min, respectively. The aux gas temperature was set to 350 °C, and the maximum spray current was established at 100.00 µA. The data collection for MS detection utilized the parallel reaction monitoring (PRM) technique, where the specific conditions and transitions for adenosine were 268.104 m/z (precursor ion) and 136.062 m/z (product ion). The analyte’s retention time was recorded as 2.83 minutes. The acquired MS spectra were processed using the Xcalibur® software from Thermo Fisher Scientific.

### Generation of co-culture spheroids

To generate co-culture spheroids consisting of 10% of MSTO-211H and 30% of Jurkat (T-cells line), we pre-staining cells with CMFDA (Cell Tracker Green CMFDA; Invitrogen) and PKH26 (PKH26 Red Fluorescent CellLi nker Kit; Sigma–Aldrich) respectively according to manufacturer’s instructions. Jurkat T cells were stimulated using phorbol 12-myristate 13-acetate (PMA) and ionomycin (Cell Stimulation Cocktail 500X, ThermoFisher Scientific) while MSTO-211H cells in the presence or absence of adenosine monophosphate (AMP). MCSs were generated as described above in 96 multi-well plates and treated with the CD73 inhibitor AB680 (Cat. No.: HY-125286, MedChemExpress). Optical images of co-culture spheroids were acquired with Opera Phenix High-content screening system (PerkinElmer) after 7 days measuring cell viability with the “CellTiter-Glo® 3D Assay” kit (Promega). Co-culture spheroids were subsequently collected, and the expression levels of IL-2, IFN-γ, and CD69 mRNA were quantified by qRT-PCR. T cell activation was assessed by measuring CD69, an early activation marker of T cells, and IL-2, a signature cytokine of activated T cells widely used as a biomarker for monitoring T cell activation (**ref PMID: 35212542**).

### Protein Quantification Analysis

Protein quantification was performed using Progenesis QI for Proteomics v4.2 (Nonlinear Dynamics) with a label-free approach. Raw data were imported and aligned based on ion intensity maps, accepting alignment scores above 60%. Peak detection used default sensitivity, with a peak width of 0.15 min and charge states +2, +3, and +4. Quantification relied on survey scan data without MS/MS, normalized across all proteins. Protein identification was done via *Mascot*, and abundance was calculated using the sum of all unique peptide normalized ion abundances for that protein in each run ^39^. Proteins showing a fold change >1.5 and p ≤ 0.05 (ANOVA) were considered statistically significant.

### Orthotopic model of mesothelioma

The study was approved by the Italian Ministry of Health (Project number B15F2.12). It was performed on immunocompetent BALB/C mice after the intraperitoneal (i.p) injection of 10^^5^ AB1-B/c-LUC cells (sarcomatoid phenotype, ^41^) using In Vivo Imaging System (Perkin Elmer). Three days after inoculation, the mice were randomly separated into six experimental groups (n=5). Drugs were administered via i.p. injection, and individual group were treated with Anti-Mouse PD-1 Antibody (RMP1-14_aPD-1) at a dose of 4,4 mg/kg, A2Br inhibitor (TT-4) and CD73 inhibitor (AB680) 3mg/kg alone/or in combination with aPD-1. The mice in the control group were treated with drug vehicle. Drugs were injected three times a week for 3 weeks. Tumor volumes were monitored by IVIS Imaging (Revvity). Each mouse received an i.p. injection of luciferin/kg body weight (IVISbriteTM D-Luciferin Potassium Salt Bioluminescent Substrate_Revvity). During image acquisition, animals were maintained under gas anesthesia (2-3% isoflurane and 1 lt/min O2). Signal intensity was quantified using Living Image Software 4.1 (PerkinElmer).

### Histopathological Examination on Mice Tissue

Formalin-fixed and paraffin-embedded tumor samples were cut into 4 μm thick sections and stained with hematoxylin and eosin for morphology. The degree of inflammation was semi-quantitatively scored according to the ratio between the severity of inflammatory infiltrate and the area examined, as follows: 0 (absent), 1 (mild), 2 (moderate), and 3 (severe inflammation). For the severity of the inflammation, the ratio was estimated by observing at least 10 fields at 40× magnification per animal. Images were captured using an Aperio slide scanner and analysed using ImageJ software v1.50i (National Institutes of Health, Bethesda, MD) ^42^.

### Immunohistochemistry on Mice Tissue

Immunohistochemical staining for evaluating inflammatory infiltrate was performed using a well-established protocol described elsewhere. Briefly, 4 μm thick sections of liver were placed on positively charged glass slides (Bio-Optica, Milan, Italy). For anti-gen retrieval, a pretreatment was created using a heat-induced epitope retrieval (HIER) citrate buffer pH 6.0 (Bio-Optica, Milan, Italy) for 20 min at 98 ⍰C. For tumor immunohistochemistry, the following primary antibodies were employed: anti-CD3 epsilon antibody [SP7], rabbit monoclonal (Abcam, ab16669); anti-FOXP3 antibody, rabbit monoclonal (Abcam, ab215206); anti-CD45 antibody, mouse monoclonal (Abcam, ab10558); and anti-PD-L1 antibody (Abcam, ab279292). Antibodies were used at the manufacturer-recommended dilutions.

Following this, endogenous peroxidase (EP) activity was blocked by applying 3% hydrogen peroxide (H2O2) for 15 min at room temperature, and then the sections were incubated for 30 min with protein blocking solution (Biocare Medical LLC). The primary antibodies were diluted in phosphate-buffered saline (0.01 M PBS, pH 7.2), added to the slides and incubated overnight at 4 ⍰C. Horseradish peroxidase (HRP) polymer was added for 30 min at room temperature and antigen–antibody reaction was visualized using the 3,3⍰-diaminobenzidine (DAB) chromogen diluted in DAB substrate buffer. Finally, the slides were counterstained with hematoxylin. Between all incubation steps, slides were washed two times (5 min each) in PBS. To test the specificity of staining and according to the most recent and relevant guidelines, in the corresponding negative control sections, the primary antibody was either omitted or replaced with an irrelevant and unspecific IgG. The inflammatory cell phenotypes were determined according to the staining pattern of antibodies against cell surface proteins. The results of immunohistochemical staining were evaluated semi-quantitatively by counting the number of immunolabeled cells in 10 fields randomly selected with a light microscope at x400 magnification.

### Shotgun proteomics

Spheroid samples derived from MSTO-211H cells were lysed with 0.2% RapiGest SF (Waters, MA, USA) with 50 mM ammonium bicarbonate on ice for 2 h, denatured at 80 °C for 15 min and then sonicated. After centrifugation (14,000 rpm, 30 min at 4 °C), protein content in supernatant was quantified by the Bradford assay. A total of 10 µg of protein was reduced (10 mM DTT), then alkylated (24 mM iodoacetamide, 37 °C for 1 h) (Sigma Aldrich, Merck KGaA) and digested with trypsin (1:50 w/w) (Promega). The samples were desalted with a C18 resin tip (Millipore, Merck KGaA), dried in a vacuum system and approximately 5 µg of peptide was resuspended in 0.1% trifluoroacetic acid and injected into a Dionex UltiMate 3000 nanosystem (Thermo Fischer Scientific) coupled with an AmaZon ETD mass spectrometer (Bruker Daltonics). Peptides were loaded onto a Pepmap precolumn (2 cm × 100 μm, 5 μm), followed by separation on a 25 cm nanocolumn (0.075 μm, Acclaim PepMap100, C18, Thermo Fischer Scientific) at a flow rate of 300 nL/min. Multistep 360-min gradients of ACN were used. The mass spectrometer equipped with a nanoBoosterCaptiveSpray™ ESI source was operated in data-dependent acquisition mode. For MS generation, enhanced resolution and a trap ICC value of 400,000 were used; for MS/MS acquisition, the ICC target was increased to 1,000,000. CID MS/MS fragmentation was set to the twenty most abundant MS peaks (top 20). The obtained chromatograms were generated using Compass Data Analysis™ v.4.2 (Bruker Daltonics) and the resulting mass lists were processed with the Mascot search engine (v.2.7.0) and searched against the human SwissProt database ^43^. Trypsin as an enzyme, carbamidomethyl (C) as a fixed modification and oxidation (M) as a variable modification were set as the search parameters. The mass tolerances for all identifications were generally fixed at 2 Da for the precursor ions and 0.8 Da for the product ions. Proteins were identified with a global FDR⍰<⍰5%, and at least one unique identical peptide sequence (p value⍰<⍰0.05) ^44^.

### Protein Network Analysis

Differentially expressed proteins were analyzed using *Ingenuity Pathway Analysis* (IPA) (GeneGo Inc), which integrates curated protein interaction and metabolic data from the literature. Networks were visualized as nodes (proteins) connected by edges (functional relationships). To complement this, STRING (Functional Protein Association Networks) was used to reveal high-confidence protein–protein interactions ^40^, further linking key regulatory proteins to major oncogenic pathways and biological processes.

## Results

### CD73 and A2Br expression are inversely related to overall survival of patients with Me

Given their possible immunosuppressive role in Me, we explored the possible predictive and prognostic role of *ADORA2B* and CD73 (*nt5E*) for immunotherapy efficacy. We first downloaded the survival data of the 82 TCGA-MESO cohort from GEPIA2 (Gene Expression Profiling Interactive Analysis) ^45^ divided respectively into 41 high and 41 low expressing specimens for each of the analyzed targets. In particular, comparing high and low *ADORA2B* or *nt5E* expression. Kaplan-Meier analysis revealed that patients with higher *ADORA2B* or *nt5E* expression had worse overall survival. The Log-rank test unraveled a significant difference between the two groups (*p*= 0.0063 and *p*= 0.012 respectively **figure 1a, 1d**). The retrospective analysis of mRNA expression of *ADORA2B* and *nt5E* of our case series (64 ME samples compared to 7 normal samples) revealed a highly significant increase in expression of *nt5E* in tumor samples (*p*<0.0042 **; **figure 1b, 1e**). In particular significantly higher *nt5E* expression was detectable in non-epithelioid subtype (** *p*< 0.01, **figure 1f**). On the other hand, we detected no significant differences in the *ADORA2B* expression in Me patients even when stratified between epithelioid (blue dots) and non-epithelioid subtypes (red dots) when compared to each other or control patients (**figure 1c)**.

**Figure 1.**
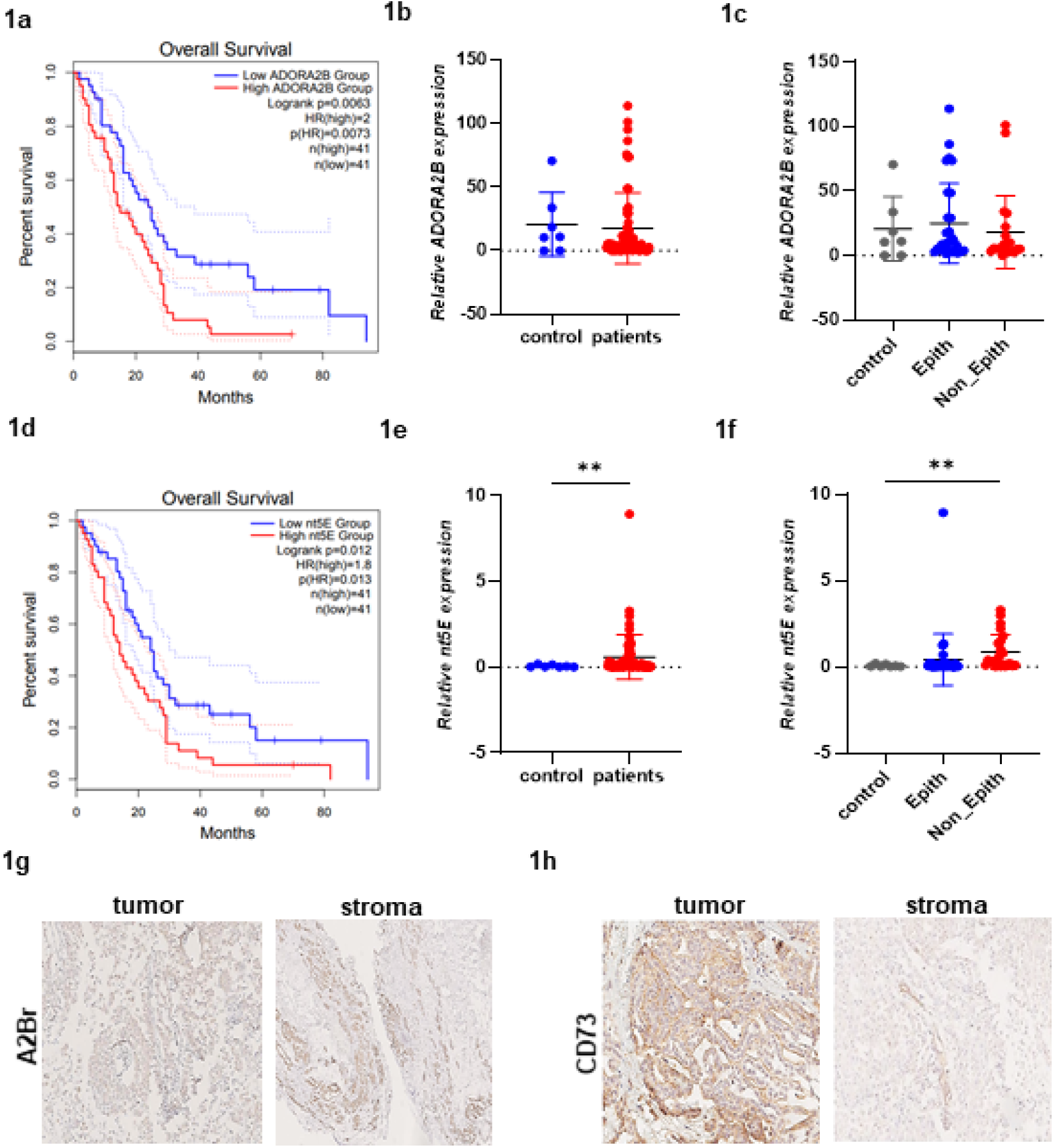
(**1a, 1d**) The Kaplan-Meier curve clearly show that low expression levels of *ADORA2B* receptor and *nt5E* (or CD73) are inversely related to better overall survival (OS) in the cohort of patients with Me tumor tissue (TCGA_GEPIA). (**1b, 1e**) *ADORA2B* and *nt5E* expression were investigated in a cohort of Me specimen compared to normal mesothelium by real-time qRT-PCR and normalized to β-actin expression. We found a differential expression in tumors compared to controls (left histogram) with *p<*0.0042 for *nt5E* expression in this cohort (** *p*< 0.01, Paired t test). (**1c and 1f**) Analysis of the possible correlation between the targets and the epithelioid (blue dots) or non-epithelioid histotype (red dots), reveal a significant correlation between *nt5E* and the latter histotype compared to the controls (Dunn’s multiple comparison test: ** *p*< 0.01). IHC analysis shows that A2B receptor and CD73 ecto-nucleotidase were expressed on tumor MM tissue form patients. Representative images of tumor-stroma expression for both are shown (**1g-1h**).

We then performed a Kaplan-Meier analysis of OS in our patient cohort, revealing a median of 12 ± 1.5 months, consisted with reported data for advanced-stage ME. Chemotherapy showed a strong positive association with OS (**Suppl. Figure 1a**, *** *p*<0.001). We then correlated adenosine markers expression with patient outcomes. Notably, CD73 high expression was significantly associated with poorer survival outcomes (**Suppl. Figure 1b**, ** *p*<0.01) while *ADORA2B* high expression showed a trend toward reduced OS (p=0.093) (**Suppl. Figure 1c**).

To validate the expression and clinical significance of the same biomarkers at protein level, we performed an immuno-histochemical analysis scoring the expression of A2Br and CD73 in the tumor cells and in the surrounding stroma (**figure 1g, 1h**). Coherent with the results of the mRNA detection, we found higher expression of A2Br and CD73 in tumor tissues. Our results show higher expression of *ADORA2B* and CD73 in tumor cells (in particular in non-epithelial cells) when compared to normal mesothelial cells. CD73 high expression showed was significantly correlated with worse patient survival (p<0.013, data not shown).

In summary, all these results confirm that although the evidenced differences in expression between epithelioid and non-epithelioid Me and the need of a broader study population, A2Br and CD73 can be considered markers of poorer survival for Me patients.

### Hypoxic conditions increase expression of CD73 and A2Br in Me cells

A panel of human Me cell lines was analyzed by Western blotting to assess the expression levels of the adenosine receptors in comparison to NP2 mesothelial cells (**Suppl. Figure 2a, 2b**). The mRNA expression profile of these markers in a set of Me cell lines, alongside four normal controls (Met5A, LP9, NP1, and NP2) is presented in **Suppl. Figure 2c**. Densitometric analysis (**Suppl. Figures 2d and 2e**) confirmed a marked increased expression of both A2Br and CD73 in Me cells compared to the NP2 normal cell line. Given the link between adenosine production and hypoxic environment ^46^, we recreated hypoxic conditions by treating Me cells with cobalt chloride (CoCl2) ^47^. To confirm if this surrogate method induced hypoxic conditions, we demonstrated activation of HIF-1α (**Figure 2**). Both *ADORA2B* and *nt5E* mRNA and protein levels were significantly upregulated across all subtypes following 6 hours of CoCl_2_ treatment. Specifically, a significant upregulation of mRNA expression was observed in REN (**Figure 2a**) and MSTO-211H (**Figure 2c**) cells, with *p*< 0.001 and *p*< 0.05, respectively. Therefore, our results show that hypoxic conditions exert a positive regulatory control of CD73 and A2Br in Me cells.

**Figure 2.**
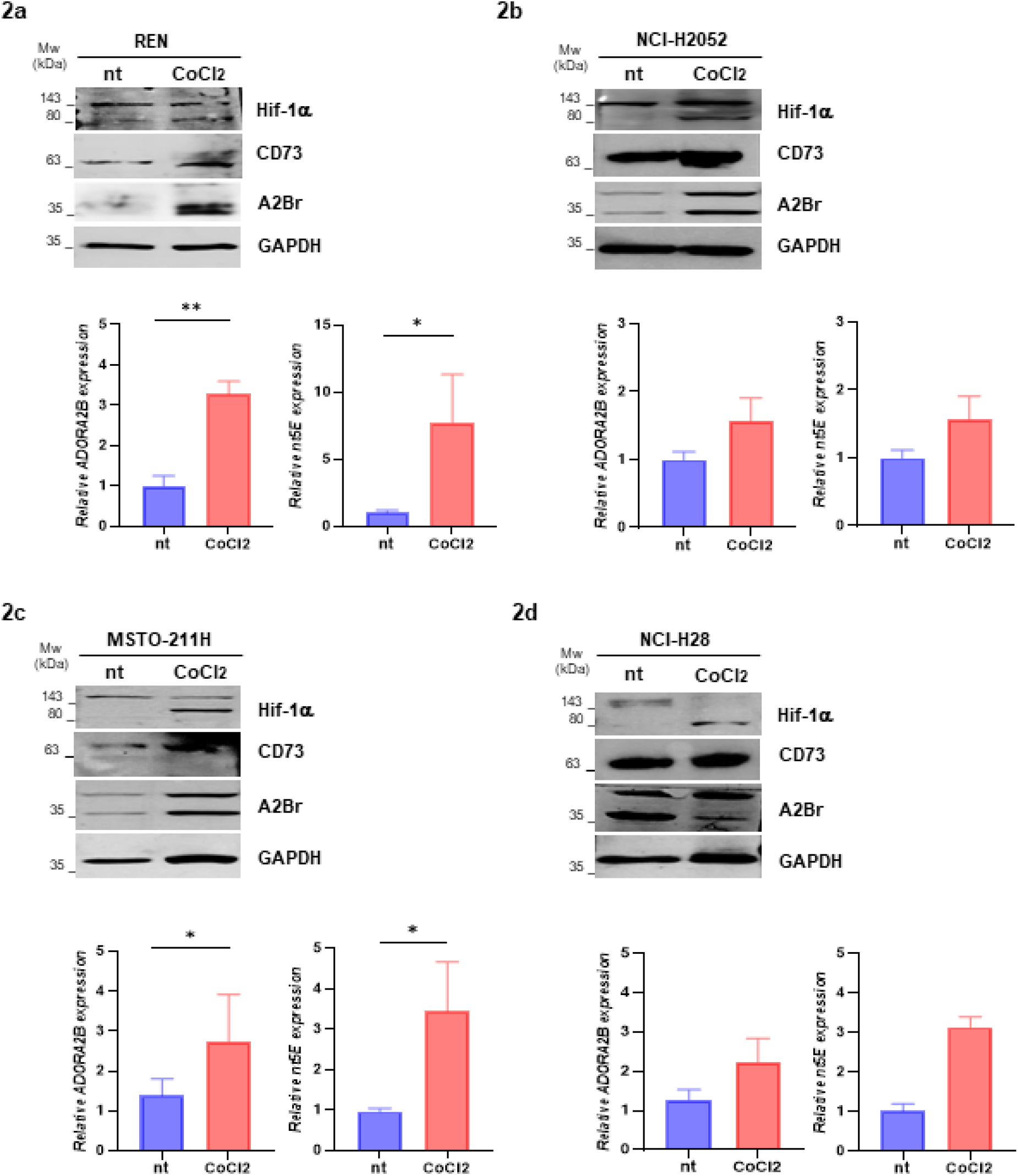
Human Me cell lines representing mesothelioma histologies: REN and NCI-H2052 (epithelioid), MSTO-211H (biphasic) and NCI-H28 (sarcomatoid) cells, were incubated for 6 h with CoCl_2_ (chemical mimic of hypoxia) to determine the expression of adenosinergic pathway by Western blotting and real time qRT-PCR. (**2a-d**) R epresentative blots were shown on the left; HIF-1α was used as a redout of hypoxia induction activation while GAPDH served as a loading control. Histograms report *ADORA2B* and *nt5E* relative expression (mean ± SD) from three independent analyses in the four Me lines. P values were calculated in REN and MSTO-211H by paired two-tailed Student t tests (* *p*<0.05, ** *p*< 0.01; n=3).

### Me cells impair immune response via adenosine production in 3D models

To assess the ability of Me cells to produce adenosine in hypoxic conditions we used a multicellular spheroids as a 3D culture model. We initially found a marked increase of *nt5E* expression (*p<* 0.05), along with elevated levels of *ADORA2B*, in 3D spheroid cultures compared to conventional 2D monolayers (**Figure 3a**). To further investigate the role of CD73 in adenosine production, we silenced its expression using specific siRNA in MSTO-211H (**Figure 3b**) and REN cells (**Supplementary Figure 3a**). Efficient transfection was confirmed in **Figure 3c** and **Supplementary Figure 3a**.

**Figure 3.**
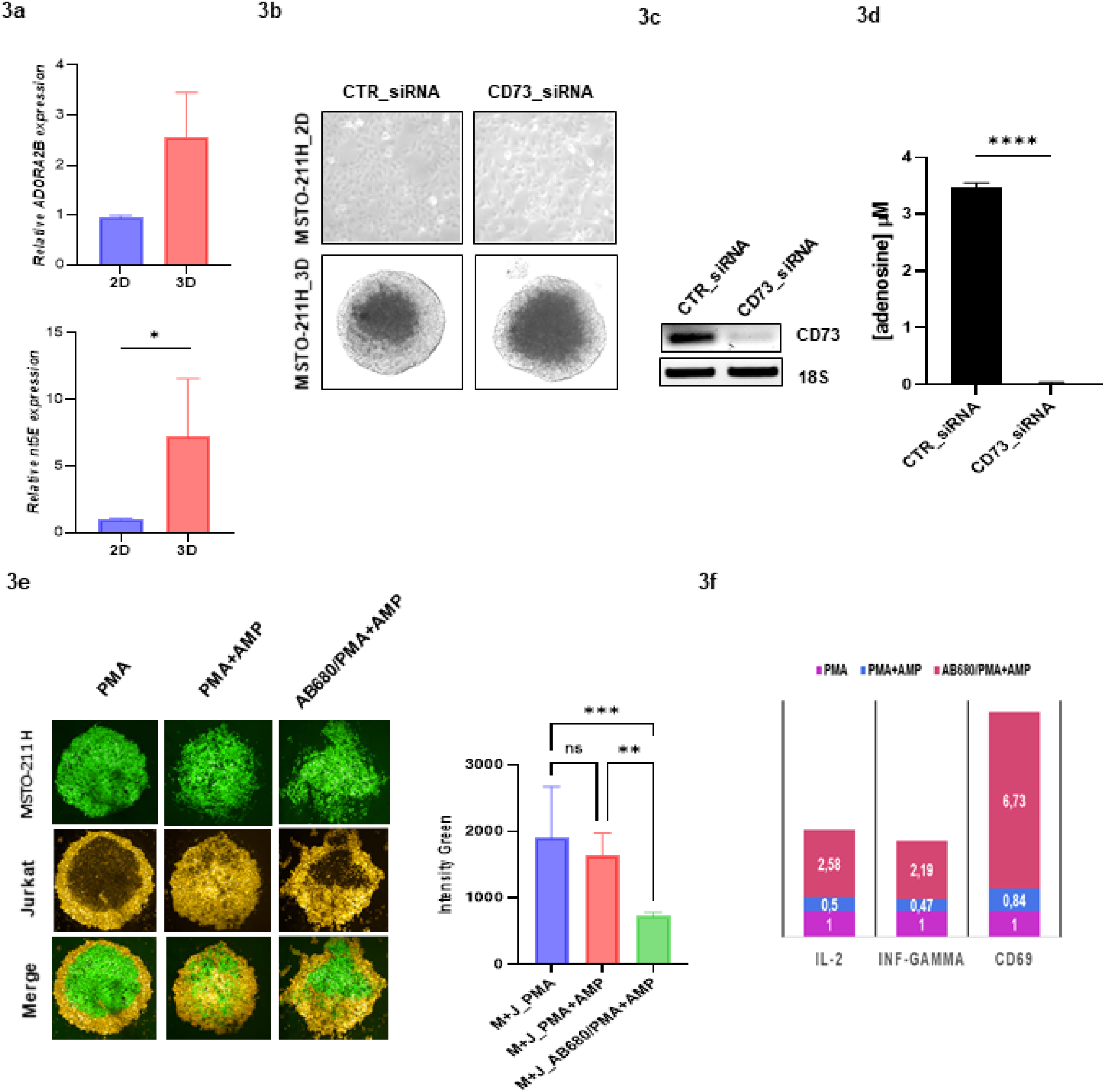
Graphs represent the relative expression of *ADORA2B* and *nt5E* (2D *vs* 3D, **3a**). MSTO-211H cells were transfected with CD73-siRNA or control siRNA and cultured in 2D and 3D conditions to evaluate the impact of CD73 silencing on cell growth. Representative images of both models are shown on the lef (**3b**). To assess transfection efficiency, *nt5E* band densities were normalized to their respective 18S bands (**3c**). Adenosine levels in 3D cultures were measured using a semiquantitative assay (**3d**), with statistical analysis performed by Student’s t-test (*p< .05) for *nt5E* expression; ****p< 0.0001 for adenosine levels; n = 3). Co-culture spheroids of MSTO-211H (green cells) plus Jurkat T cells (yellow) were stimulated with PMA and Ionomycin cocktail, in the presence or absence of adenosine monophosphate (AMP) and in combination with AB680 (iCD73) as treatment. Here, we reported the images acquired at Opera Phenix (PerkinElmer) with the quantitative analysis of fluorescence intensity in the graph (**3e**). Two-way ANOVA was used to evaluate statistically significant differences (*** *p*<0.0001, ** *p*< 0.01; n = 3). Expression of genes coding for IL-2 and IFN-γ and CD69, as an indicator of T cell activation, were measured by qRT-PCR. (**3f**).

Semiquantitative analysis of extracellular adenosine levels in 3D cultures revealed a significant reduction following *nt5E* silencing (**Figure 3d** and **Supplementary Figure 3b**, **** *p*<0.0001). This provides the evidence that Me cells can generate adenosine via CD73, and that this capability could influence the immune response in the tumor microenvironment. Therefore, we investigated the immunomodulatory role of CD73 using the AB680 inhibitor (CD73i) in co-culture models of MSTO-211H (green cells) and CD73-negative Jurkat T cells (yellow cells). Jurkat cells were stimulated with PMA/Ionomycin and co-cultured in spheroids in the presence or absence of the ATP-precursor AMP and/or AB680. Fluorescence intensity, measured using Opera Phenix imaging, revealed that CD73 inhibition significantly affected MSTO-211H cell viability in co-cultures system (**Figure 3e**, *** *p*<0.001). Treatment with AB680, markedly enhanced immune activation within the co-culture spheroids, as indicated by increased IFN-γ expression and the upregulation of the T cell activation markers IL-2 and CD69 (**Figure 3f**).

### Antagonists of the Adenosine pathway exert a significant immune-mediated reduction on murine Me growth

A2B receptor and CD73 expression was confirmed by Western blot analysis in the murine Me cell lines representing the three major histotypes: AB1 AB22, and AB12 (**Figure 4a**). Based on these results, we developed an orthotopic ME model in immunocompetent BALB/c mice as shown in **Figure 4b**. Immunocompetent BALB/c mice were intraperitoneally injected with 10^^5^ AB1-B/c-LUC cells. Three days post-injection, to assess the effect on tumor growth, mice were randomized into six treatment groups and received therapy every three days for three weeks as follows: PBS (control=ctr), TT-4 (a selective A2Br antagonist; Ki = 9.1⍰nM; 3⍰mg/kg), AB680 (CD73 inhibitor; 3⍰mg/kg) and their combinations with aPD-1 therapy (Clone RPM1-14; 4,4⍰mg/kg). Tumor burden was assessed by bioluminescence imaging (IVIS) after luciferin injection, and signal intensity quantified with Living Image Software. Representative images from three mice per group show tumor changes from T0 to T21 of treatment (**Figure 4c**). Single agent treatment TT-4 or AB680 exerted significant reductions in tumor volume *vs* control (**** p< 0.0001) while combined treatments with aPD-1 plus either TT4 or AB680 resulted in a greater reduction in tumor growth *vs* single agents alone (**** p< 0.0001). In particular treatment with aPD-1 alone did not lead to a significant reduction in tumor growth compared to control (**Figure 4d**). Body weight was monitored throughout the study as an indicator of treatment-related toxicity (**Figure 4e**).

**Figure 4.**
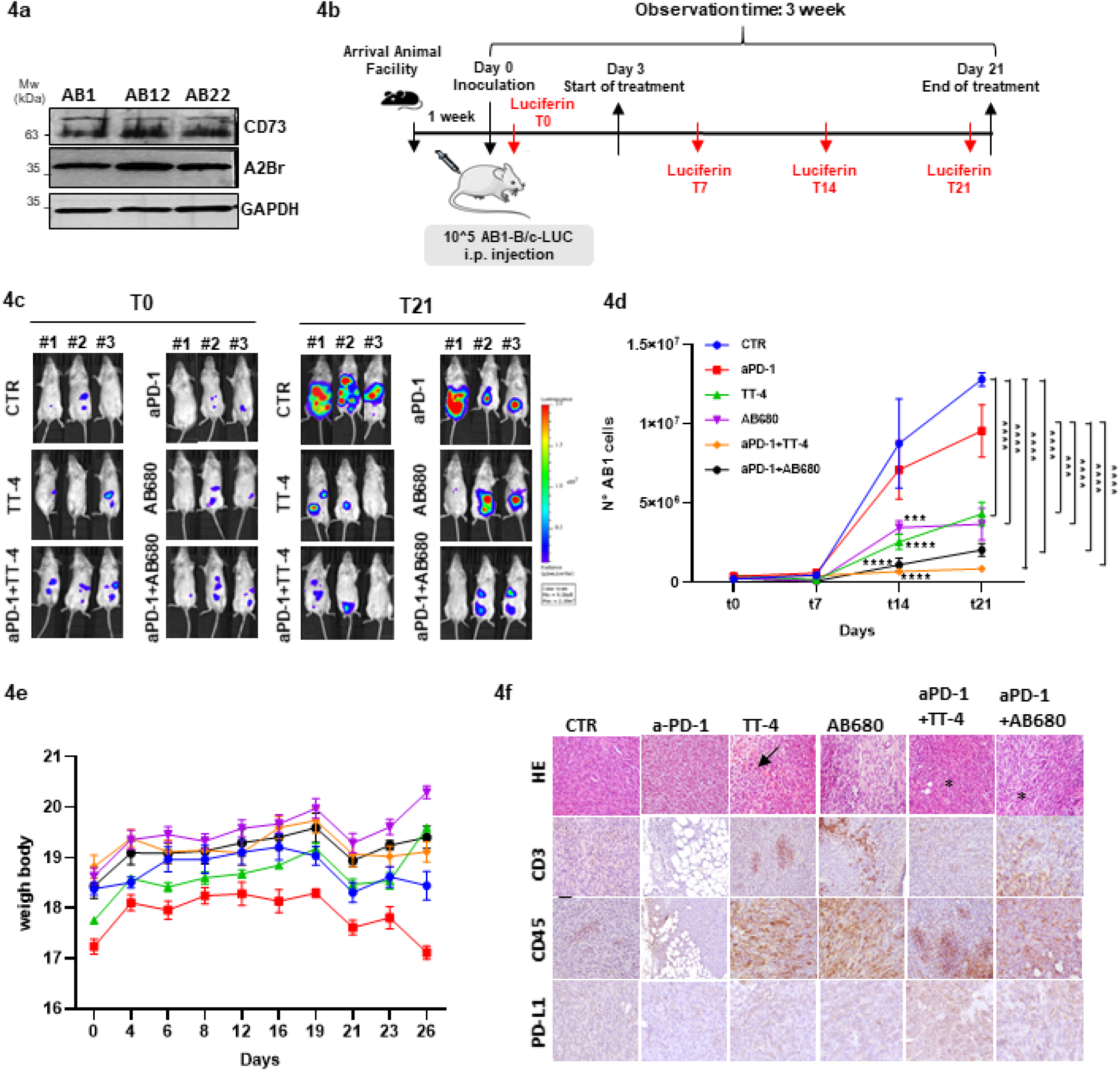
Western blot analyses confirmed the expression of A2Br and CD73 in murine Me cell lines: AB1 (sarcomatoid), AB22 (biphasic) and AB12 (epithelioid). Band densities were normalized against their respective GAPDH (**4a**). A schematic representation of the program followed during the development of the orthotopic model of the Me was reported (**4b**). Immunocompetent BALB/C mice were inoculated with 10^5 AB1-B/c-LUC cells i.p.----Three days after, mice were randomized into six groups and treated q3days × 3 weeks: 1) PBS 100 μl (control= ctr); 2) aPD-1 (4.4 mg/Kg); 3) TT-4 (a clinic ready, selective A2Bri, Ki = 9.1nM, Portage Biotech, 3 mg/Kg); 4) AB680 (CD73i, 3 mg/Kg); 5) TT-4 plus aPD-1; 6) AB680 plus aPD-1. Tumor volumes were monitored over 21 days via IVIS bioluminescence imaging following luciferin administration. Signal intensity (ROI) was quantified using Living Image Software 4.1 (Revvity). Representative bioluminescence images of three mice/group at the start (T0) and end of treatment (T21) are shown, normalized to the fluorescence scale on the right (**4c**). The curves report the mean ± standard error of the mean (SEM) of individual tumor volumes of mice at the indicated time point (n=5). Statistical differences were assessed by two-way ANOVA with Tukey’s multiple comparisons test (** *p*< 0.01, **** *p<*0.0001, **4d**). Body was tracked throughout the study as an indicator of treatment-related toxicity and is presented as means ± SEM in the graph (**4e**). Formalin-fixed, paraffin-embedded tumor samples were sectioned (41μm), stained with hematoxylin and eosin for morphological assessment, and evaluated under a light microscope (40× magnification). Stained sections were semi-quantitatively evaluated for the percentage and localization of positive cells, focusing on CD3^+^, CD45^+^ and PD-L1 markers (**4f**). Across all groups, neoplastic cells displayed typical mesothelioma morphology-intertwined bundles with mild to moderate extracellular matrix and moderate mitotic activity (2–3/HPF). Inflammatory infiltrate was minimal in the control and aPD-1-only groups. In contrast, tumors from the TT-4 and AB680 groups exhibited moderate to severe lymphoplasmacytic infiltration (arrows), with no significant differences between them. Combination treatments (TT-4 + aPD-1 and AB680 + aPD-1) showed multifocal, chronic lymphoplasmacytic infiltrates often localized around necrotic areas (asterisks), again with no significant difference between these two groups. Immunohistochemical analysis confirmed T and B cell infiltration (CD3+ and CD45+), with CD45 expression significantly higher in all treatment groups compared to control. PD-L1 was expressed in neoplastic cells across all groups compared to control and aPD-1 monotherapy groups.

In the aPD-1 treatment group, mice showed variable responses: some exhibited transient weight gain likely due to ascites and an underlying inflammatory response, while others experienced weight loss associated with diarrhea, suggesting gastrointestinal side effects.

Overall, these findings reflect a mixed toxicity profile with signs of localized inflammation rather than severe systemic toxicity. Histopathological analysis was conducted on formalin-fixed, paraffin-embedded tumor tissues (4 µm) using hematoxylin and eosin staining. Immunohistochemistry was performed for CD3^+^, CD45^+^, CD4^+^, Foxp3^+^, CD20^+^ and anti-PD-L1.

Sections were scored semi-quantitatively for the percentage of positive cells and/or their localization. Images were acquired using a 40× objective lens on a light microscope (**Figure 4f**). Morphological analysis revealed consistent tumor architecture across all groups, characterized by interwoven cellular bundles, moderate extracellular matrix, and low mitotic activity (2–3/HPF). Inflammatory infiltrates were minimal in both the control and aPD-1 groups.

Although CD45^+^ cells were predominant following aPD-1 treatment, the scarce presence of CD3^+^ cells, absence of CD4^+^, and rare FOXP3^+^ cells reflected to limited activation of the adaptive immune response, likely due to an immunosuppressive tumor microenvironment. In contrast, TT-4 and AB680 monotherapies induced moderate to severe lymphoplasmacytic infiltration. Notably, combination therapies resulted in moderate, multifocal inflammation centered on necrotic areas and significantly increased immune infiltration compared to monotherapies and control (* *p*< 0.05 and ** *p*< 0.001, respectively). This was associated with organized, non-encapsulated lymphoid aggregates rich in CD20^+^ B cells surrounding high endothelial venules (HEVs), indicative of tertiary lymphoid structures (TLS) with germinal centers. Concurrently, tumor cells displayed moderate and diffuse membranous and cytoplasmic PD-L1 expression that neither changed after treatment compared with controls nor changed with the response (**Supplementary figure 5a-b**). The addition of TT-4 or AB680 to aPD-1 significantly enhanced immune cell infiltration (CD45^+^/CD20^+^), indicating a shift toward a more immunogenic and responsive tumor microenvironment than that obtained with aPD-1 only.

These findings support the rationale for combining immune checkpoint blockade with adenosine pathway inhibition to boost antitumor immunity and improve therapeutic outcomes in primary or secondary ICIs resistant tumors.

### Proteomic analysis after treatment of Me cells with adenosine agonist shows a deep effect on a broad range of oncogenic and immune-suppressive pathways

Label-free proteomic analysis was applied to identify and quantify proteins whose abundances significantly differed between untreated and NECA or A2Br antagonist MRS1754 treated MSTO-211H-derived spheroids. Progenesis software was used to quantify the differentially expressed proteins using the following filters: fold change⍰≥⍰1.5 and ANOVA test p value⍰≤⍰0.05. In details, we identified 905 differentially expressed proteins modulated by NECA treatment (308 upregulated and 525 downregulated) and 1258 modulated by the A2B antagonist (768 upregulated and 490 downregulated, respectively). The pie chart in **Supplementary figure 6a** illustrates the distribution of proteins modulated by each treatment individually. A principal component analysis (PCA) plot derived from unsupervised multivariate analysis of differential protein expression profiles demonstrated close correlations between the spot maps of the biological replicates, thus highlighting the experimental reproducibility (**Figure 5a-b**). To enhance the translational relevance of A2Br antagonist-mediated immunomodulation, a correlation network was generated to explore potential interactions between A2Br and Key cancer-associated networks. Pathway enrichment analysis of the 93 proteins commonly affected by both treatments was further conducted using Ingenuity Pathway Analysis (IPA), focusing on direct interactions (**Figure 5c**, Additional file 1). This analysis identified TP53, MYC, STAT1, and JUN as central regulatory nodes (highlighted in green). These proteins are functionally interconnected and play pivotal roles in key biological processes, including inflammatory signaling, T cell development, and modulation of the tumor microenvironment (also highlighted in green, right), critical aspects of tumor progression and immune evasion. These proteins were found to be part of a single, high-confidence interaction network, functionally associated with key biological processes such as cellular development, cell proliferation, and essential molecular functions (represented in light blue, yellow, and pink, left panel). In addition, miR-27 (colored in green) emerges as a critical regulatory hub, known to drive tumor progression by modulating multiple oncogenic signaling pathways: including Ras/MAPK, AKT, TGF-β, and Wnt/β-catenin (highlighted in light blue and yellow, at the left of network) ^48^, ultimately influencing cancer cell proliferation, survival mechanisms, and immune evasion. YAP1 (highlighted in orange at the bottom of the network) was found to be upregulated following NECA treatment, confirming previous observations. This was paralleled by the identification of the 14-3-3 pathway (in light blue on the left side of the network), which is known to modulate YAP/TAZ activity.

**Figure 5.**
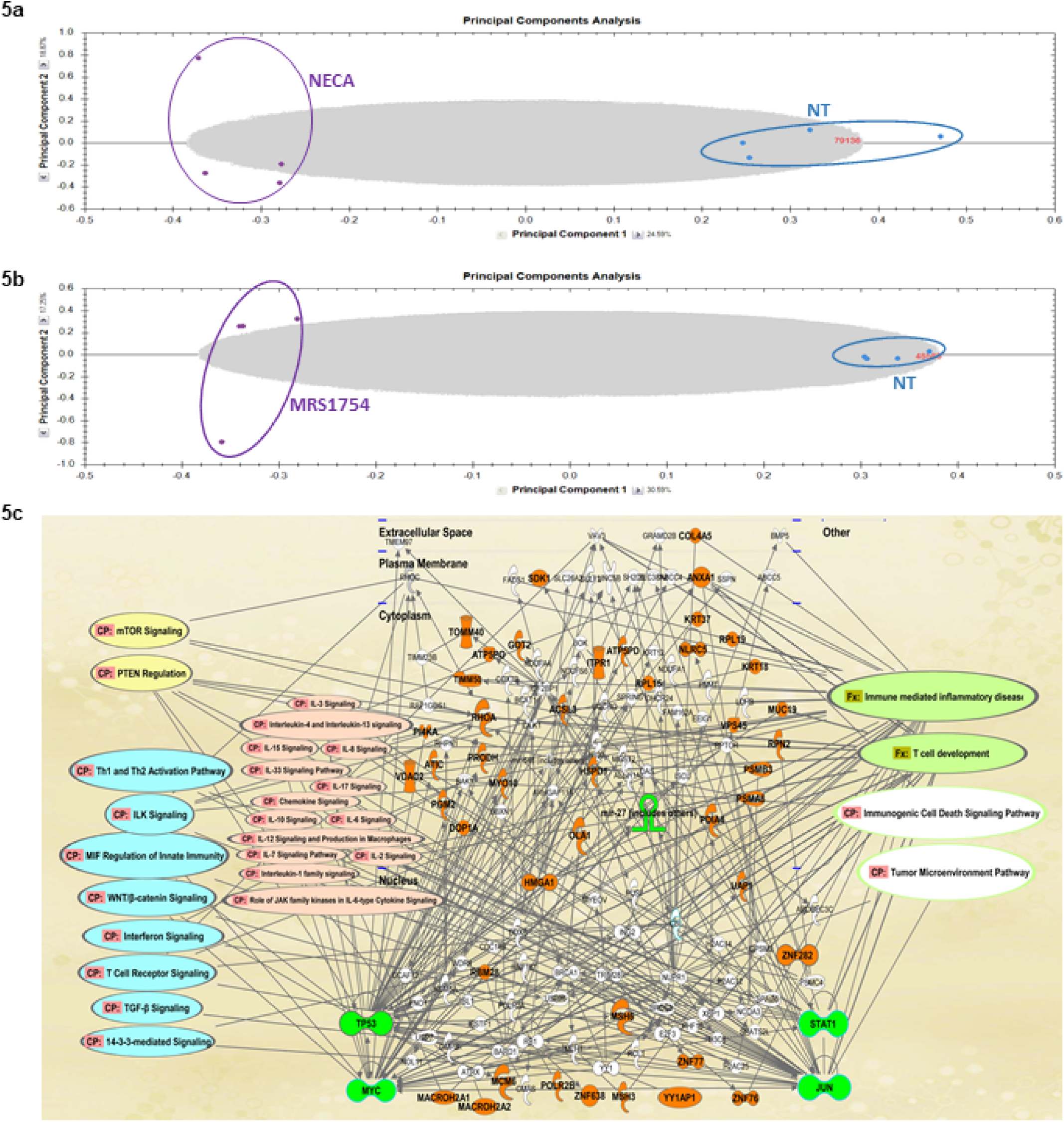
Unsupervised multivariate analysis using PCA of differential protein expression profiles revealed strong clustering of biological replicates, confirming both the robustness and reproducibility of the experimental approach (**5a-5b**). A network of differentially expressed proteins was identified in MSTO-211H-derived spheroids treated for 5 days with 1001nM NECA and 0.71nM of the A2Br antagonist MRS1754. IPA of 93 commonly modulated proteins revealed TP53, MYC, STAT1, and JUN as central nodes (colored in green), associated with key pathways involved in inflammation, T cell development, and the tumor microenvironment (also highlighted in green, right) (**5c**). Proteomic analysis of treated MSTO-211H spheroids, with 1001nM NECA and 0.71nM of the A2Br antagonist MRS1754, identified 93 shared differentially expressed proteins. IPA revealed TP53, MYC, STAT1, and JUN as central nodes (colored in green) linked to inflammation, T cell development, and the tumor microenvironment (also highlighted in green, right). Among these, 46 statistically significant proteins form a single, high-confidence interaction network (score 7.0) associated with cellular development, proliferation, and key molecular functions (colored light blue, yellow, and pink, left). MiR-27 (colored in green), emerges as a regulatory hub that promotes tumor progression by modulating key signaling pathways, including Ras/MAPK, AKT, TGF-β, and Wnt/β-catenin influencing cellular proliferation, survival, and immune response.

STRING analysis supports the IPA-based network of the 93 modulated proteins by confirming key interactions among central regulators and oncogenic pathways (**Supplementary figure 6b**). These results delineate functionally interconnected pathways implicated in tumor progression, thereby facilitating the identification and validation of therapeutic targets. Pharmacological modulation with NECA and its antagonist MRS1754 was instrumental in uncovering deregulated proteins, although further investigations are required to define the specific mechanistic roles of individual candidates. Collectively, the data support a model in which adenosine promotes immunosuppression within the tumor microenvironment through receptor-mediated signaling, operating via both autocrine and paracrine mechanisms

## Discussion

The current study highlights a prognostic role of the adenosine pathway in Me and provides an experimental model to prove the therapeutic effect of its inhibition. We demonstrate that, although to a lesser extent, the expression levels of both CD73 and A2Br expression are associated to a poorer overall survival of Me patients and that hypoxia induces their over-expression. Moreover, we demonstrate that Me generate adenosine via CD73 and that in 3D model CD73 inhibition promotes immune response. Our murine syngeneic model confirms how hampering adenosine production or the effects of adenosine on Me cells respectively exerts a potent immune-mediated anti-cancer effect via CD73 and A2Br inhibition. Eventually we provide evidence that the Adenosine analogue NECA deeply influences the proteomic pattern of Me cells toward an oncogenic and immunosuppressive profile.

Under inflammatory and hypoxic conditions, activation of the adenosinergic pathway acts as a negative feedback mechanism to limit excessive tissue damage resulting from sustained immune cell activation and adenosine levels in the tumor microenvironment are 10 to 20 times higher than those in healthy tissues ^80^, indicating the presence of immunosuppressive concentrations of adenosine in tumors ^80^. Therefore a role for the adenosine pathway in human tumors is increasingly attracting attention ^34, 36, 53-56^. In renal cell carcinoma, liver cancer, colon cancer and pancreatic adenocarcinoma (PDAC) there is evidence supporting the notion that the inhibition of the adenosine pathway may represent a strategy to modulate the immune milieu and improve therapy response ^37, 57-60^ whereas in Triple Negative Breast Cancer (TNBC) and NSCLC increased CD73 levels are associated with poorer prognosis ^61-63^. Nevertheless, there still a need to derive a better understanding of the relevance of this pathway in cancer to improve immune responses in most human tumors. To our knowledge, the present study provides the first body of evidence supporting the adenosine pathway as a new actionable target to boost immune response in Me cells and overcome the weak and unpredictable response of this tumor to ICIs ^17^.

In this report we have demonstrated on both Me tissue and cell lines that Me tumor cells express two pivotal components of the adenosine pathway namely the ectonucleotidase CD73 and the A2Br involved in adenosine production and adenosine-dependent signaling respectively. The negative prognostic impact of tumor expression of CD73 and A2Br, observed in patients with Me is consistent with the existing mesothelioma data set as exemplified by our analysis of the TCGA dataset using GEPIA2.

Hypoxia is a key factor hampering resistance to ICi, chemotherapy and radiotherapy ^64-66^, and adenosine production is promoted by hypoxia ^36^. Given that Me is an hypoxic tumor ^35, 67^ we simulated hypoxic conditions in ME cell lines using the HIF inducer cobalt chloride (CoCl2) to assess its effect on both A2Br and CD73. After 6 hours, both the mRNA and protein expression levels of A2B and CD73 were significantly increased in REN and MSTO-211H cells, with increased phospho-CREB suggesting functional activation of the A2B receptor under hypoxic conditions (Costa *et al*., data not shown).

These results provide the first direct evidence that HIF-1α-mediated hypoxia drives transcriptional regulation of CD73 and A2Br in Me cells. To further explore the effects of the adenosine pathway and, in particular, of CD73 in Me cells we employed both 2D monolayer and 3D spheroid culture models derived from MSTO-211H and REN cell lines before and after *NT5E* silencing. The experiments clearly demonstrated that CD73 knockdown reduces ME cells proliferation, via adenosine/A2Br binding (data not shown) This effect was even more pronounced in the 3D spheroid models, which better recapitulate the in vivo tumor architecture and gradients of oxygen and nutrients. Notably, the silencing of CD73 led to a marked expansion of the hypoxic core within spheroids, a finding that suggests that CD73 may contribute to maintaining a viable proliferative niche under oxygen-deprived conditions. Although the pro-proliferative effect of the adenosine pathway on tumor and Me cells is not new ^68, 69^ our results highlight the role of CD73 to maintain a viable proliferative niche under oxygen-deprived conditions.

Moreover, we have demonstrated that Me cells produce adenosine via CD73 and, through a functional co-culture assay of Me cells treated with the CD73 inhibitor AB680 and CD73-negative Jurkat T cells^71^, we showed that Me cells not only release adenosine into the TME but also exert a direct adenosine-dependent immunosuppressive effect.

The inhibition of CD73 has previously been shown to promote T cell activation and proliferation, deplete Tregs and support the maturation and functional restoration of natural killer (NK) cells ^55^ and while this is not the first demonstration of adenosine production by tumor cells ^70^, we provide the first evidence of an immune-suppressive role of CD73 in Me. Threfore CD73 expression by these cells is not just a prognostic biomarker but, on the contrary, it exerts an active rife intra-tumor immune-suppression that facilitates tumor immune evasion by shaping the biochemical milieu ans should be taken into account prior to considering immunotherapy for Me. The CD73 inhibitor AB680 led to a marked enhancement of T cell activation and function as demonstrated by a significant increase in the cells picked from the coculture inflammatory cytokines IFN-γ and T-specific IL-2 (REF)., two key mediators of anti-tumor immune response The reactivation of T cell effector was further confirmed by the concomitant significant upregulation of CD69, an early marker of T cell activation. These results therefore provide solid evidence that reversing adenosine production activates T cell response and reinforces the notion that the adenosine pathway is a key-regulator of tumor immune suppression within TME.

The results obtained with the co-cultures prompted us to test an orthotopic ME mouse model in immunocompetent BALB/c mice using luciferase-expressing CD73 and A2Br positive AB1 MM murine cells. Treatment with either an A2B antagonist or a CD73 inhibitor led to a marked immune response along with reduction of the tumor growth, underscoring the strong antitumor potential of the adenosine pathway blockade and providing evidence of a pivotal role of A2B as an actionable therapeutic target

Notably, combining these agents with anti-PD-1 further enhanced therapeutic efficacy, whereas on the contrary anti-PD-1 alone had minimal impact. These findings highlight the lack of clinical efficacy of PD-1 inibition in Me ^13^ and support the rationale for combining adenosine inhibition with other ICIs to better counteract the immunosuppressive TME. Histopathological analyses of the murine tumors revealed that anti-PD-1 alone failed to induce substantial T cell infiltration while monotherapies with TT-4 or AB680 only moderately increased immune cell presence. Critically, combination treatments significantly enhanced immune infiltration and promoted the formation of organized tertiary lymphoid structures (TLS), characterized by CD45^+^ and CD20^+^ cells, germinal centers, and high endothelial venules. Although the presence of TLS and stromal B cells are considered a common predictive biomarker of response to ICI ^72^ our results show how the very treatment with adenosine and anti-PD-1 elicits such a strong immune response to give origin to TLS. These features are indicative of a robust and coordinated antitumor immune response and support the therapeutic integration of CD73 and A2Br inhibitors for Me particularly in combination with anti-PD-L1 immune checkpoint inhibitors.

To deepen the translational potential of A2Br antagonism in Me, we conducted an in-depth proteomic analysis following treatment of MSTO-211H-derived 3D spheroids with NECA and the antagonist MRS1754. Mass spectrometry identified broad alterations in the tumor proteome: NECA modulated 905 proteins, while A2Br inhibitor MRS1754 affected 1.258, suggesting a significant impact of A2Br signaling on cellular function.

Importantly the same analysis unraveled that 93 proteins were commonly influenced by both treatments, whereas ingenuity pathway analysis (IPA) revealed that oncogenic regulators: TP53, MYC, STAT1, and JUN played a pivotal role within this network. Given the key role played by these genes in inflammation, T cell development, and tumor microenvironment modulation these results highlight the critical immunological relevance of the A2B axis. Furthermore, 46 proteins formed a statistically significant and highly interconnected network associated with core biological processes such as cell growth, differentiation, and immune regulation. One notable finding was the identification of miR-27a as a critical regulatory hub. This miRNA is, known to activate multiple oncogenic pathways (RAS/MAPK, AKT, TGF-β, Wnt/β-catenin) ^48^, all of which contribute to tumor progression and immune escape. Moreover, the study linked A2B signaling with the Hippo-YAP/TAZ pathway, a central axis in Me pathobiology due to frequent NF2 mutations ^51, 73^ and its frequent aberrant activation thay plays a pivotal role in tumor promotion, immune evasion and therapy resistance of this tumor.

Elevated YAP1 contributes the establishment of an immunosuppressive TME by promoting the expression of inhibitory mediators such as PD-L1, IL-10, and TGF-β, suppressing effector T cell responses, and fostering the expansion of regulatory immune populations like Tregs and M2 macrophages. In this context, NECA-induced YAP1 upregulation plays a pivotal role in maintaining immune evasion and reducing the effectiveness of immunotherapies. Therefore, this signaling pathway emerges as a compelling therapeutic target, with several inhibitors currently advancing through clinical development **(REF)**. Following NECA treatment, YAP was upregulated, aligning with prior data and further supporting the involvement of adenosine signaling in therapy resistance. The detection of the 14-3-3 pathway, a known modulator of YAP/TAZ activity, further reinforces this regulatory connection. These findings emphasize the potential of A2B antagonists, alone or combined with immune checkpoint inhibitors, to disrupt tumor-promoting pathways and enhance immunotherapy efficacy.

Study limitations: This study utilizes well-established Me cell lines, and the use of primary Me cells (with matched immune cells) could certainly provide additional information supporting the role of this pathway as a therapeutic target in Me. In addition, the use of a non-epithelial Me cell line limits the *in vivo* results. However, we believe that these limitations are significantly overcome thanks to the homogenous distribution of A2Br and CD73 and by their equal sensitivity to hypoxia of both of the Me cellular subtypes we show here

It could also be argued that the lack of expression of CD39 and A2Ar in primary Me suggests that the adenosine pathway may not be fully actionable. This can be balanced by the finding of their presence in the surrounding stroma (Costa et al., data not shown). The immunohistochemistry data from the immunocompetent mouse model, which showed no significant differences in PD-L1 expression between tumors treated with single agents and control groups, supports findings from the literature indicating that, although high PD-L1 levels are associated with poor prognosis in Me, they do not effectively predict patient response to anti-PD-1 therapy ^74^.

As recently demonstrated in other tumors ^57, 75-79^, all together our data lay the groundwork for clinical strategies combining adenosine pathway inhibitors with immune checkpoint inhibitors as an approach to re-prograMe TME toward immune responsiveness, with the ultimate goal of overcome therapy resistance and improve outcomes in this challenging disease.

## ABBREVIATIONS

AOUS: Azienda Ospedaliera Universitaria Senese
ATCC: American Type Culture Collection
ADORA2A and -B receptors: A2Ar and A2Br
Me: Malignant mesothelioma
FFPE: formalin-fixed paraffin embedded
IHC: immunohistochemistry
HIF-1α: hypoxia-inducible factor 1^α^
ICIs: immune checkpoint inhibitors
PBS: phosphate-buffered saline
CoCl2: cobalt dichloride
PD-L1: Programmed death-ligand 1
qRT-PCR: real-time quantitative reverse transcription
PCR SD: standard deviation
SEM: standard error of the mean

## Declarations

## Acknowledgements

We acknowledge the support of the Gruppo Italiano Mesothelioma (https://gime.it/) and the Portage Biotech for providing the A2Bri (TT-4) for *in vivo* testing. We would like to express our sincere gratitude to Dr. Monica Cantile and Dr. Giosuè Scognamiglio of the Institutional Biobank (BBI), Pascale Institute, for providing FFPE tumor samples.

## Ethical Approval

The animal study and the retrospective analysis of human samples were conducted under current regulations and approved by the appropriate Ethics Committees, as detailed in the ‘Materials and Methods’ section.

## Consent for publication

All authors have agreed with publishing this manuscript.

## Availability of data and materials

All supporting data are available in the manuscript and Supplementary Information. Public datasets used are specified in the article. For material requests, please contact the corresponding author.

## Funding

This work was supported by research funding from the Italian Ministry of Health ‘Ricerca corrente L3/5

> ***“Patient-Derived Tumor Organoids (PDO) and Xenografts as tools for Precision Medicine”***.

## Author contributions

CC, SGG, LM GP LM contributed to the conception and design of the study; CC conducted most of the experimental work, supported by GP, LM, LA, RL, and BP; CC, RL, and FB carried out the *in vivo* studies; CB, DDB conducted the anatomo-pathology analyses; CC, RL, BP performed the bioinformatic analyses; CC, SGG, GP, LM, RL, FB, BP, CB, DDD, LM contributed to data analysis; CC, SGG, LM prepared the manuscript. All authors revised the manuscript and approved the final version.

---

**Supplementary Table S1.**
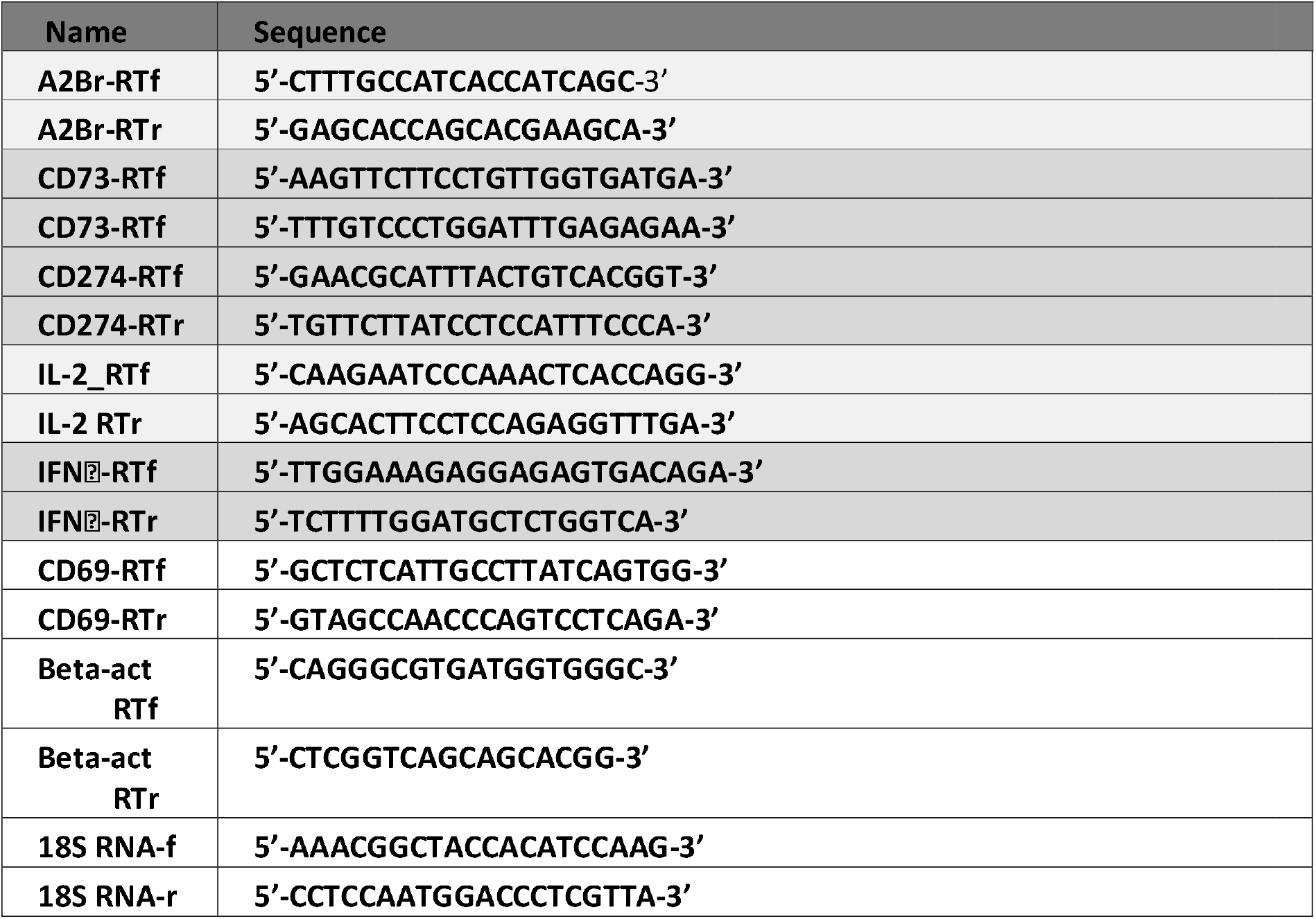

## Supplementary Figure

**Supplementary figure 1.**
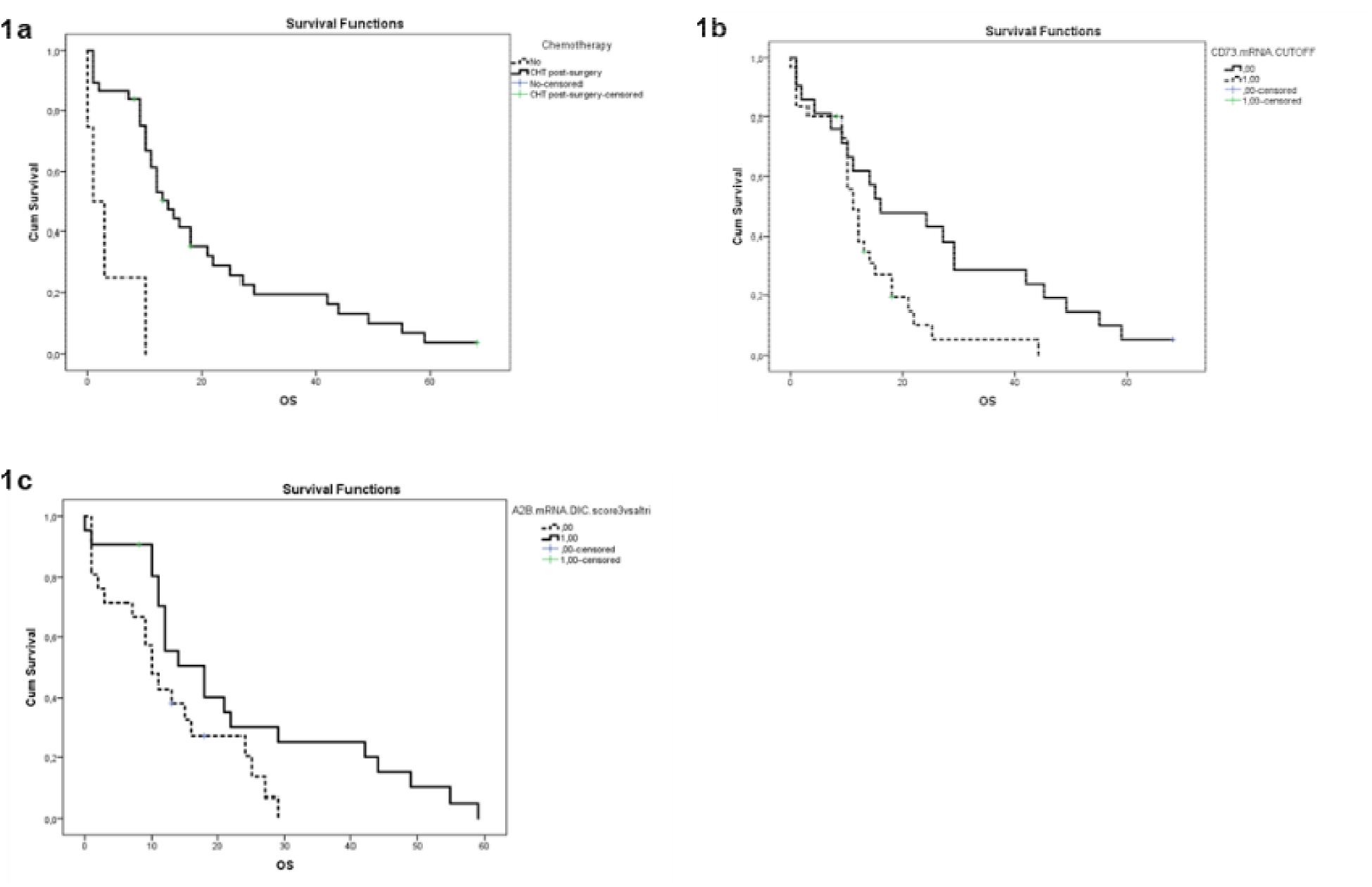
Kaplan–Meier overall survival according to type of systemic treatment: chemiotherapy versus untreated (**1a**); according to CD73 expression: low mRNA cutoff=0,00 or high mRNA cuttoff= 1,00 (**1b**); according to A2B expression: low mRNA cut off=0,00 or high mRNA cut off= 1,00 (**1c**).

**Supplementary figure 2.**
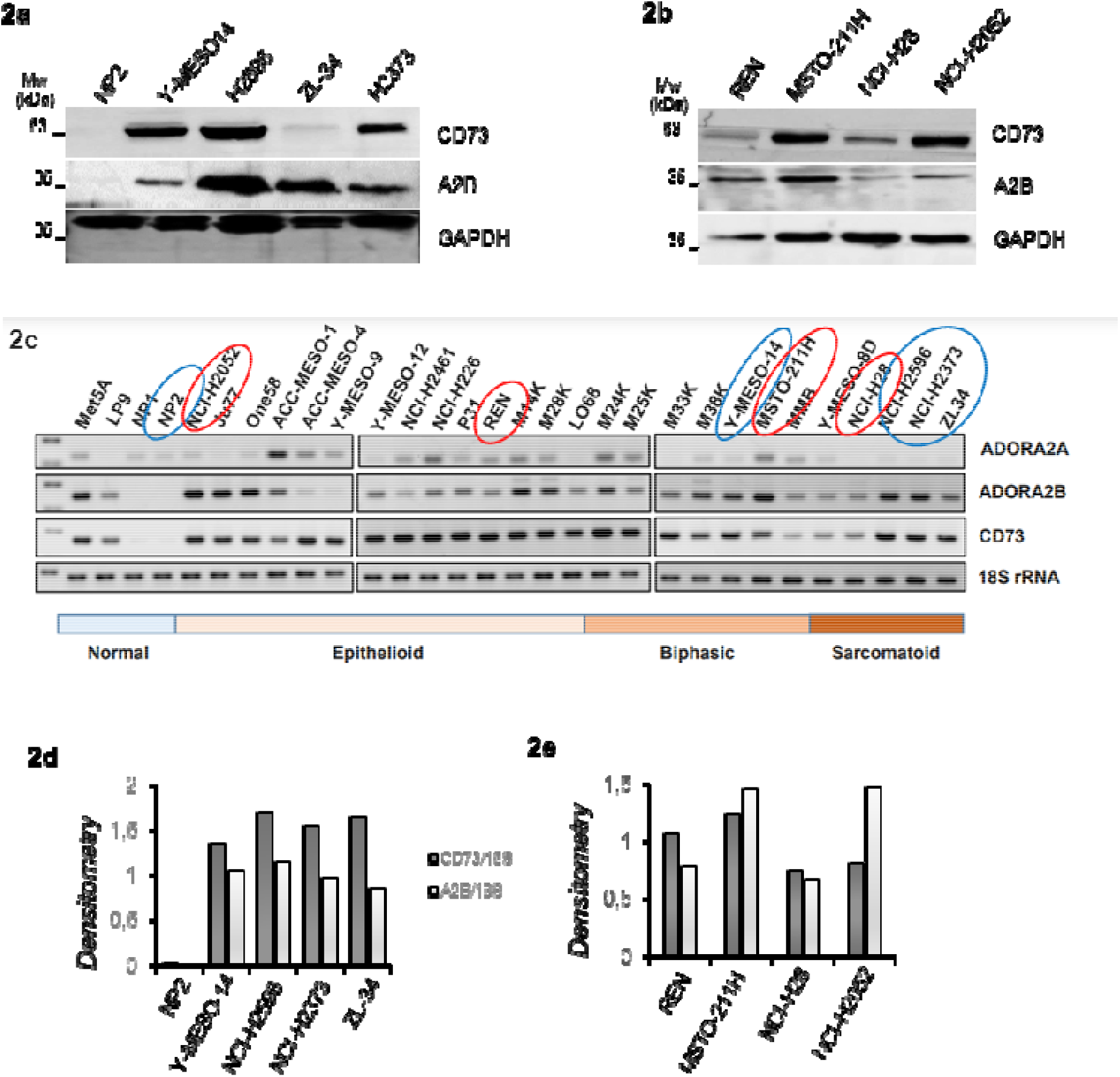
CD73 and A2B protein expression levels were evaluated in Me cell lines by Western blotting, using NP2 normal cells as a control. GAPDH was used as a loading control (**Suppl. figure 2a-b**). A panel of Me cell lines, categorized by histological subtype into epithelioid, biphasic, and sarcomatoid groups, was analyzed for CD73 and A2B mRNA expression levels and compared with four normal cell lines: Met5A, LP9, NP1, and NP2 (**Suppl. figure 2c**). Band intensities were quantified by plotting the CD73/18S and A2B/18S ratios (**Suppl. figure 2d-e**). Densitometric analysis is presented in graphs, where cell lines marked in blue are shown on the left panel, and those in red are displayed on the right panel.

**Supplementary figure 3.**
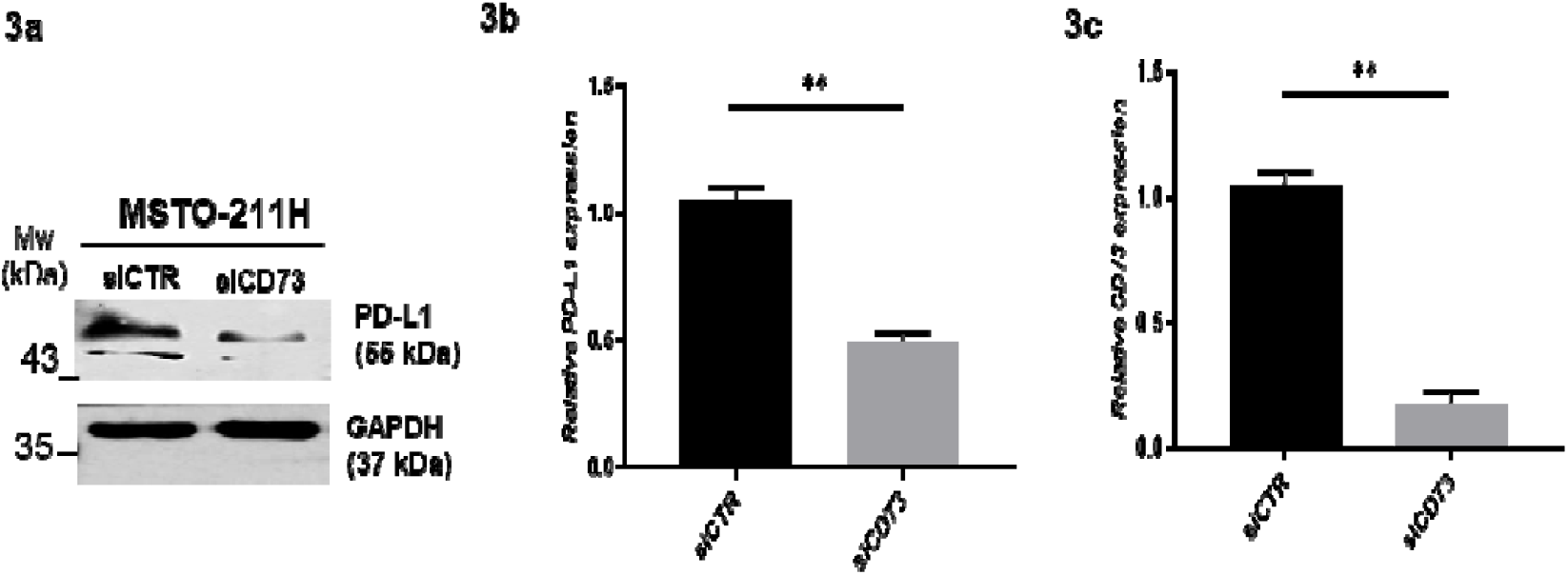
MSTO-211H cells were transfected with siCD73 and the siRNA control (siCTR). PD-L1 protein levels were analyzed by Western blotting, with GAPDH used as a loading control. A representative blot is shown (**Figure 3a**). PD-L1 mRNA expression was quantified by qRT-PCR and is presented as fold-change relative to β-actin (**Figures 3b–c**). CD73 silencing significantly reduced PD-L1 mRNA levels (** *p*< 0.01), as determined by a two-tailed t-test. The graph on the right displays transfection efficiency.

**Supplementary figure 4.**
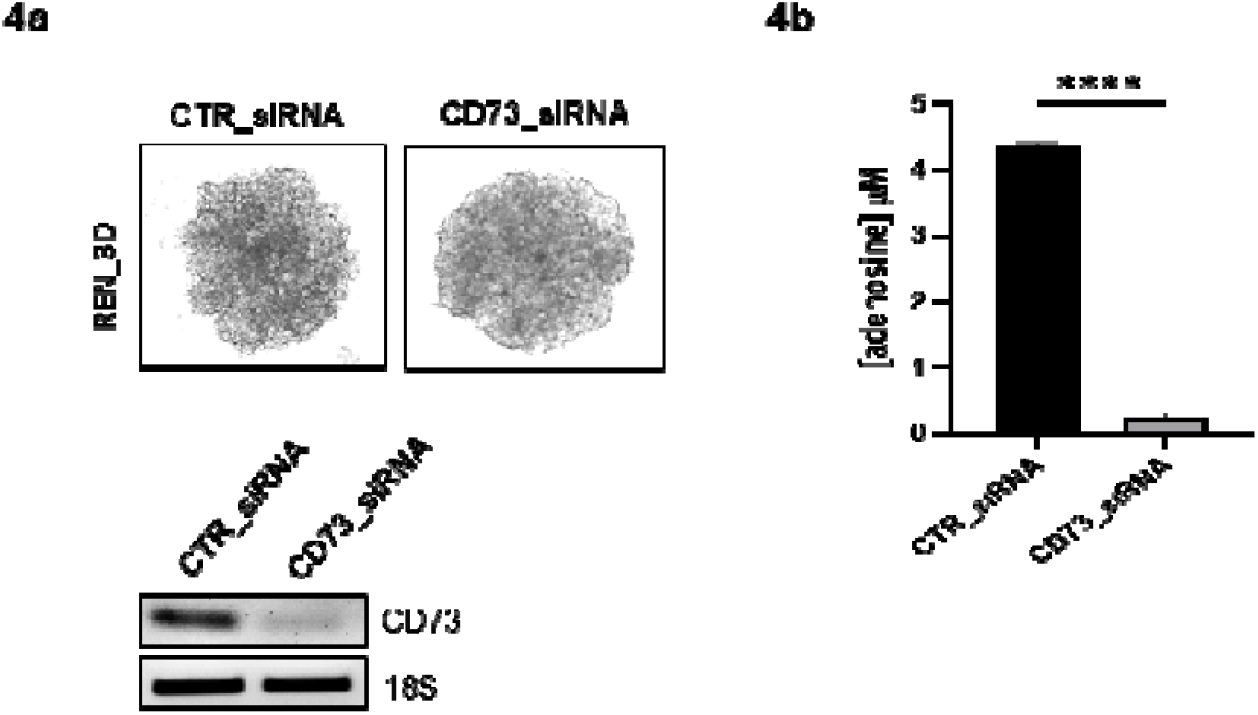
REN cells were transiently transfected with either CD73-targeting siRNA (siCD73) or control siRNA (CTR-siRNA), and extracellular adenosine levels were quantified. To generate 3D multicellular spheroids (MCS), cells were seeded in 96-well plates. Representative images of REN-MCS are shown (**Figure 4a**). The efficiency of CD73 silencing in REN-MCS was evaluated by blot analysis, with 18S serving as a loading control (**Figure 4b**). Data represent the mean ± SD from three independent experiments by two-tailed t-test (**** *p<*0.0001).

**Supplementary figure 5.**
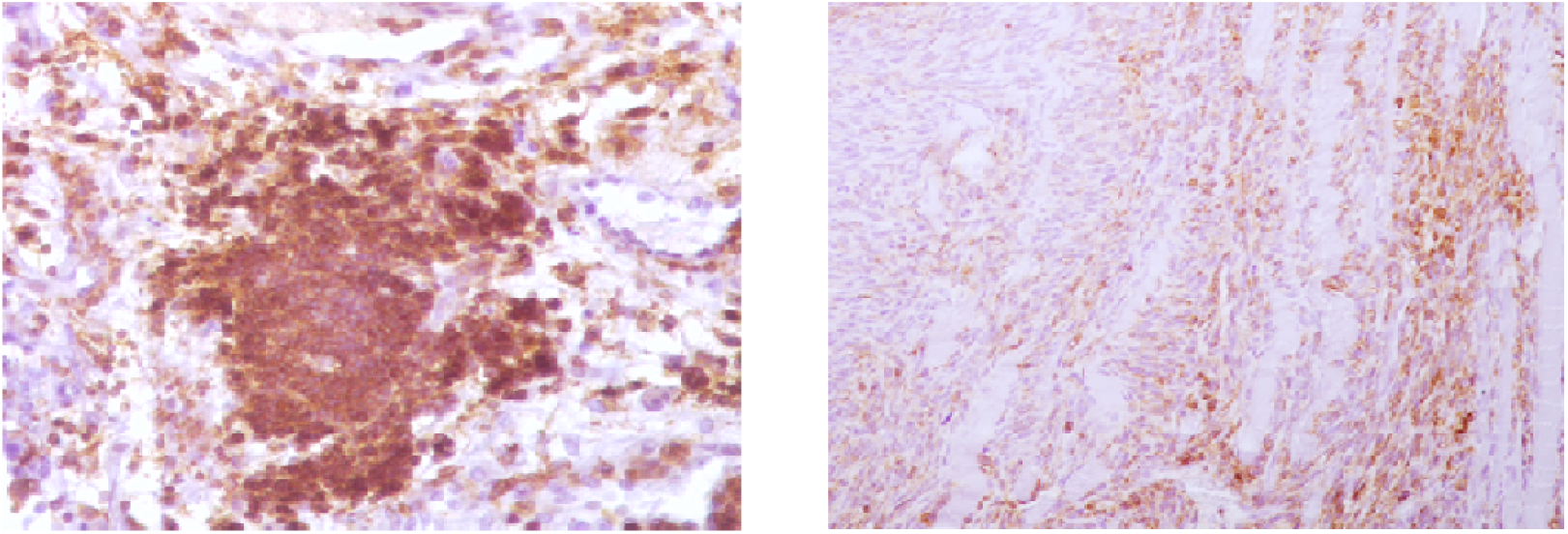
Organized, non-encapsulated, discrete and irregularly nodular aggregate of immune cells consisting mostly of CD20 + B cells around high endothelial venules (HEVs, **Figure 5a**). Neoplastic cells show scattered and moderate membranous and cytoplastic immunolabeling for Rb anti-PD-L1 antibody (**Figure 5b**).

**Supplementary figure 6.**
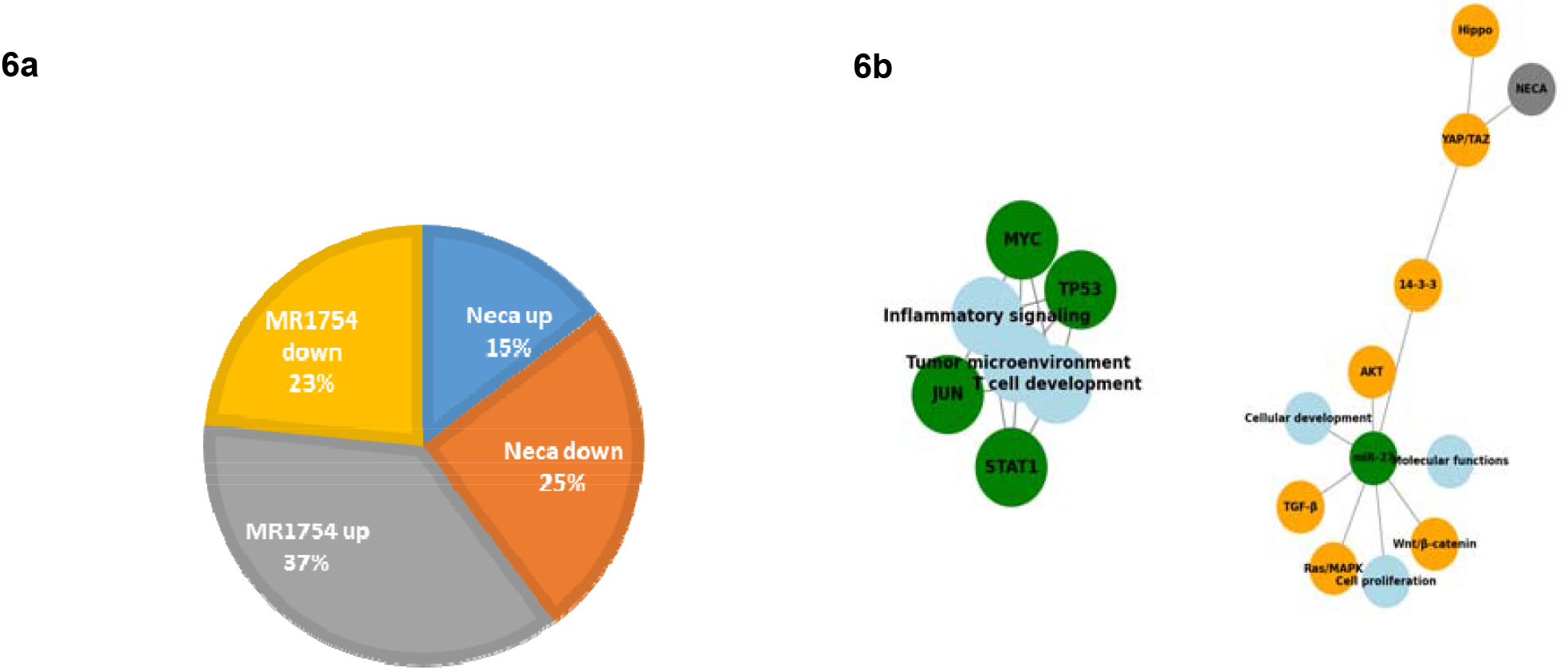
The pie chart illustrates the distinct effects of NECA and the A2B receptor antagonist on protein expression. NECA treatment resulted in 308 upregulated and 525 downregulated proteins, whereas the antagonist induced upregulation of 768 proteins and downregulation of 490 **(Figure 6a)**. STRING (Functional Protein Association Networks) complements the analysis of the 93 commonly modulated proteins by mapping high-confidence interactions between key nodes such as TP53, MYC, STAT1, JUN, and miR-27 (green) and their associated pathways. It reinforces the functional relevance of these regulators within critical biological processes (e.g., immune signaling, cell proliferation, tumor microenvironment remodeling) (blue nodes) and oncogenic pathways (e.g., Ras/MAPK, AKT, YAP/TAZ) (orange node) **(Figure 6b)**. By integrating both predicted and validated associations, STRING supports the IPA findings and helps highlight potential therapeutic targets within a coherent interaction network.

